# The genetic makeup of the electrocardiogram

**DOI:** 10.1101/648527

**Authors:** Niek Verweij, Jan-Walter Benjamins, Michael P. Morley, Yordi van de Vegte, Alexander Teumer, Teresa Trenkwalder, Wibke Reinhard, Thomas P. Cappola, Pim van der Harst

**Author notes:** Corresponding Authors: Prof. Pim van der Harst, University of Groningen, UMCG, Department of Cardiology, PO Box 30.001, 9700 RB Groningen, The Netherlands; Dr. Niek Verweij, University of Groningen, UMCG, Department of Cardiology, PO Box 30.001, 9700 RB Groningen, The Netherlands.

## Abstract

Since its original description in 1893 by Willem van Einthoven, the electrocardiogram (ECG) has been instrumental in the recognition of a wide array of cardiac disorders^1,2^. Although many electrocardiographic patterns have been well described, the underlying biology is incompletely understood. Genetic associations of particular features of the ECG have been identified by genome wide studies. This snapshot approach only provides fragmented information of the underlying genetic makeup of the ECG. Here, we follow the effecs of individual genetic variants through the complete cardiac cycle the ECG represents. We found that genetic variants have unique morphological signatures not identfied by previous analyses. By exploiting identified abberations of these morphological signatures, we show that novel genetic loci can be identified for cardiac disorders. Our results demonstrate how an integrated approach to analyse high-dimensional data can further our understanding of the ECG, adding to the earlier undertaken snapshot analyses of individual ECG components. We anticipate that our comprehensive resource will fuel *in silico* explorations of the biological mechanisms underlying cardiac traits and disorders represented on the ECG. For example, known disease causing variants can be used to identify novel morphological ECG signatures, which in turn can be utilized to prioritize genetic variants or genes for functional validation. Furthermore, the ECG plays a major role in the development of drugs, a genetic assessment of the entire ECG can drive such developments.

## Main

An enhanced understanding of the influence of genetic variants on the complete cardiac cycle represented by the ECG could generate new hypotheses on cardiac physiology, disease and effects of drugs. To better characterize the impact of genetic variants on the ECG, we obtained all 77,190 3-lead ECGs of the UK Biobank that contained raw signal data necesserary for the analysis. After individual level and population level quality control to remove abnormal beats and ECGs^3^ (**Online Methods**), 63,706 individuals remained for the primary analyses. The primary ECG morphology phenotype was constructed by dividing one representation of the averaged cardiac cycle on the ECG into 500 temporal data points, as dictated by the data sampling frequency of the stored ECG recording. To study possible effects of heart rate, we performed additional, secondary, analyses in which we normalized that representation for the individual beat-to-beat variation (the RR-interval). To demonstrate that this approach also captures the classical ECG traits, we used previously described genetic variants in aggregate and isolation to visualize their morphological effect on the ECG^4-13^. By plotting 500 association signals of each datapoint as −log_10_ P-values along the time axis of one beat (**Fig. 1** and **Supplementary Data 1**), we found that the polygenic risk score of PR-interval associated with a shift of the P-wave; the polygenetic risk score of QRS-duration associated with Q and S-wave durations; the polygenic risk score of 12-lead-sum area matched the area under the curve of the QRS complex; and the polygenetic risk score of QT-time associated with T-wave prolongation. Interestingly, when individual variants of these risk scores were plotted, a plethora of different ECG morphologies was observed, suggesting they are not bound by classic ECG patterns and indicate differences in underlying biology (**Fig. 2a-b, Supplementary Data 2**). By examining the genetic heritability along the time-axis of one heartbeat, we observed that electrically active (non-isoelectric) ECG points were more heritable and that our primary ECG phenotype (unadjusted for RR-interval) showed greater heritability (see **Fig. 2c**).

**Fig. 1.**
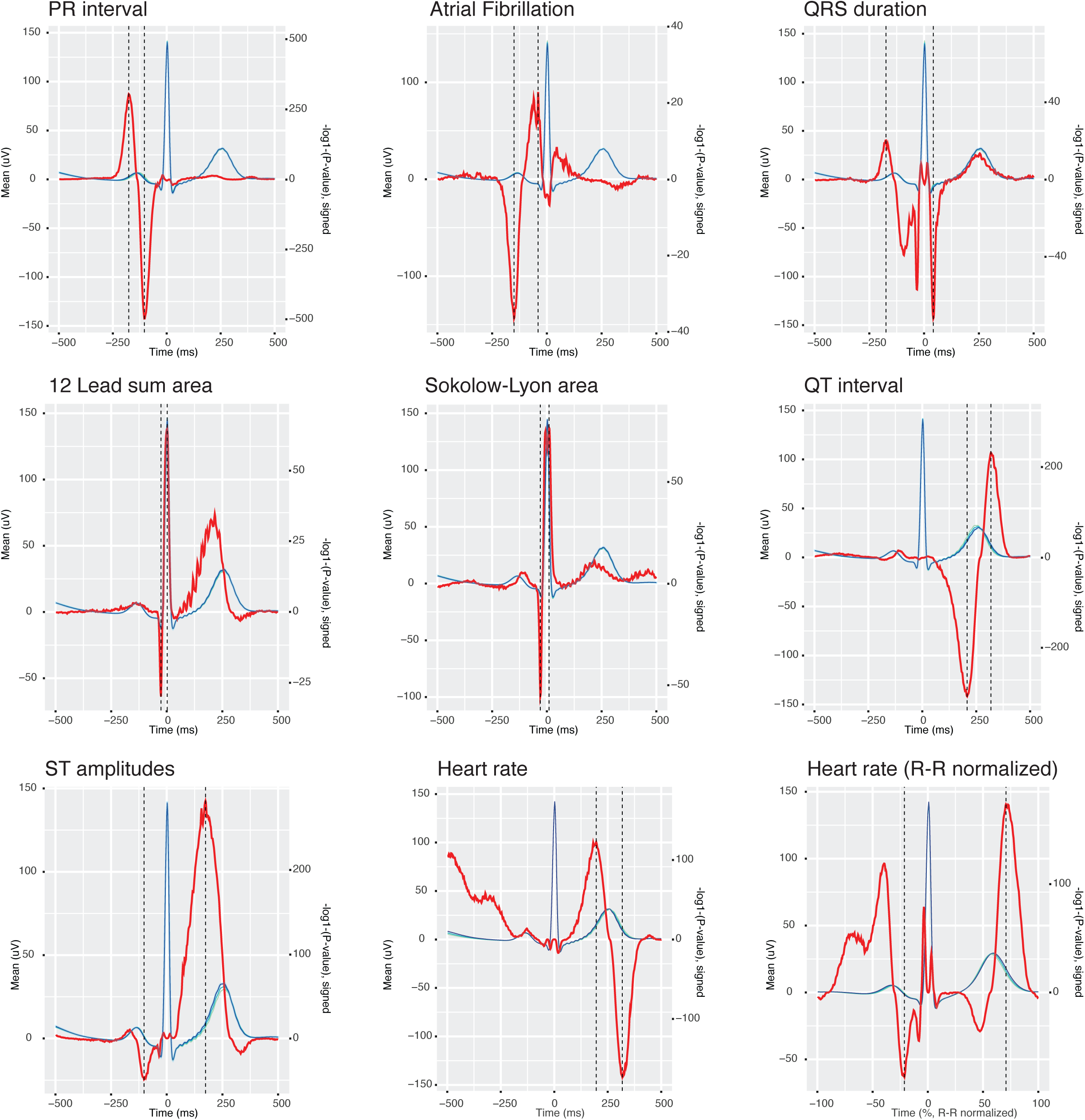
Polygenic risk scores of ECG indices are associated with the trait expected segments of the ECG. The left y-axis depicts the micro voltage scale, the y-axis on the right the signed −log10(*P*-values), the x-axis time in ms or % of RR normalized as indicated. The blue lines are the average ECG amplitude of the full cohort and the red lines the P-value for association with each time point of the ECG (n=500 timepoints) on a log_10_ scale, signed to show direction of assocation. The dashed vertical black lines mark point of strongest negative and positive association. Additional plots can be found in the supplementary information.

**Fig. 2.**
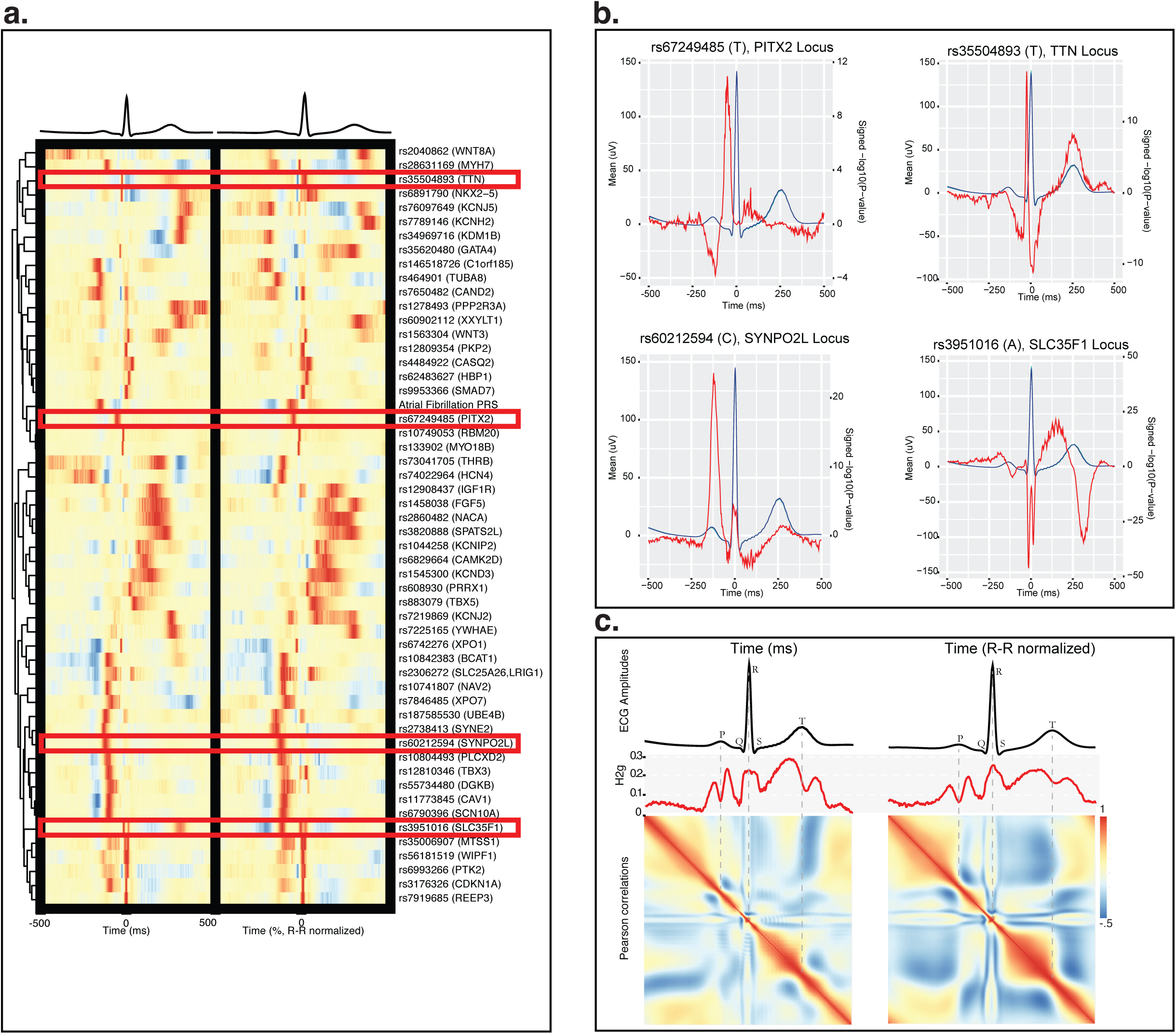
Genetic variants display unique morphological signatures and the heritability is highest at ECG segments showing greatest electrical activity. **a**, Heatmap of the normalized association patterns for genetic variants that were previously found to be associated with AF in order to compare the effects across loci. Effects were orientated to the most positively associated allele across all time points and colored in red; a blue color indicates a negative effect while yellow indicates no effect. **b.** excerpts of the impact on ECG morphology by previously reported genetic variants; the red line indicates the signed - log_10_(P-value) for association across the heart beat. **c.** The ECG-morphology phenotype of the averaged beats with and without normalization of individual beat-to-beat (R-R) intervals. The SNP-heritability’s (h^2^_g_) were highest for the ECG morphology unadjusted for R-R intervals; the maximum observed heritability was 0.29 (se=0.01) at the ST-wave segment. Heritability estimates were high for ECG segments that have high electrical activity. A Pearson correlation matrix between each ECG data point was computed across the ECG. The resulting heatmaps are presented per phenotype, with red indicating high positive correlation, blue heigh negative correlation, with yellow no correlation.

Next, we performed genome wide association studies (GWAS) on the amplitude of each of the 500 temporal data points of the ECG generating 500 association values per genetic variant (**Online Methods**). In total 414 independent (r^2^ <0.001) genetic variants were identified in 331 2MB regions at the traditional genome wide significance threshold (P<5×10^-8^) or 203 independent variants in 166 2MB regions at the stringent bonferroni corrected significance level (P<6.25×10^-10^). These 414 independent were assigned to 560 candidate genes (**Fig. 3a., Supplementary Table 1-3**). Of all 331 ECG associated regions, 127 were shared (1MB region-based) with loci found in prior GWASes of classical ECG traits; 64 regions were shared with the atrial fibrillation phenotype (**Supplementary Table 1 and 4**). We identified 179 genetic association signals that had not been reported before^4,5,14,15,6-13^. The impact of all genome wide significant genetic variants on ECG-morphology is plotted in **Supplementary Data 3,** and all genetic variants studied can be viewed online in our ECG-browser (http://www.ecgenetics.org). The three strongest observed association signals were for rs7132327 (*P*=4×10^-140^) near *TBX3*, rs6801957 (*P*=3×10^-133^) in *SCN10A* and rs2074238 (*P*=2×10^-129^) in *KCNQ1*.

**Fig. 3.**
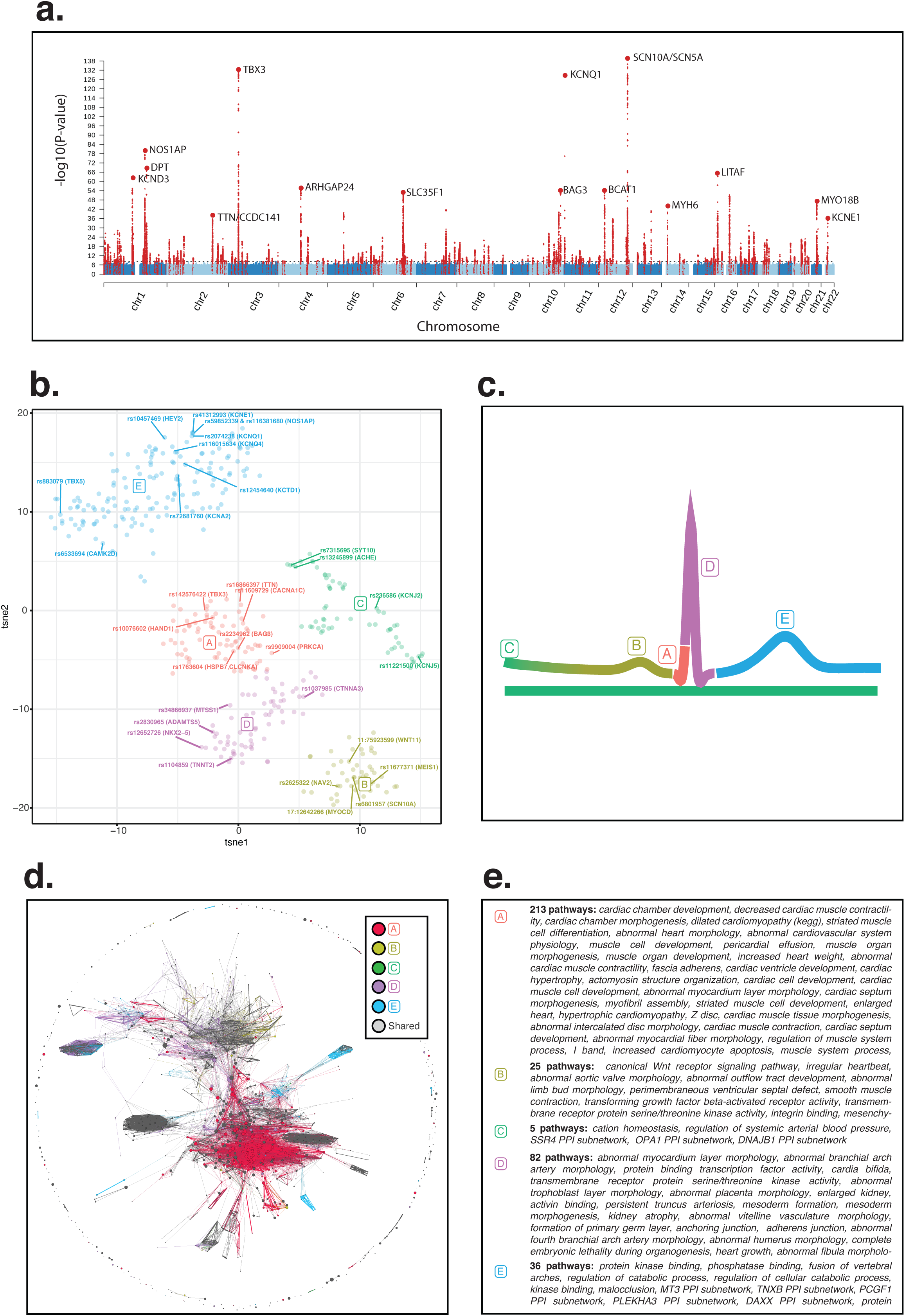
Unbiased clustering maps main effects of genetic variants on the electrocardiogram and indicate differentially enriched pathways. **a**, manhattan plot of the ECG morphology phenotype (smallest p value across all traits is shown) variants in red indicate those passing P<5×10^-8^ the most significant variants have been annotated with their nearby genes. **b**, The t-SNE plot derived from the variant-ECG morphology association patterns **c**, Illustration of how each cluster can be linked back to their primary ECG effect. **d,** Network plot of the DEPICT pathway analysis, indicating the correlation between biological gene-sets (edges), the significance (node size) and differential enrichment (colors correspond to the clusters, grey nodes indicates no differential pathway enrichment P>0.01). **e**, Differentially enriched biological gene-sets terms per cluster.

To group genetic variants with similar main effects on the ECG we performed unbiased clustering of the normalized ECG morphology association profiles. This analysis suggested 5 subsets of genetic effects on the ECG (**Fig. 3b**). Four clusters A, B, D and E contained variants that associated primarily with differences in Q-wave, P-wave, R-S Wave and T-wave morphology, respectively (**Fig. 3c**). The 5^th^ cluster (cluster C) contained a diversity of ECG morphologies, but all variants appear to influence a component of heart rate. Although genetic variants grouped into a cluster sharing morphological features, many expressed unique features within each cluster, pointing towards differences in underlying biology.

To further explore potential biological mechanisms of the ECG-associated genetic loci, we used DEPICT^16^ and identified 1,163 enriched biological gene-sets. Among the most significant gene-sets were “heart development”, “decreased cardiac muscle contractility”, and “embryonic growth retardation”. The most significant tissues identified were “heart”, “heart ventricles” and “heart atria” (**Supplementary Table 5-6**). Next, we considered the 5 distinct subsets identified by unbiased clustering analyses and explored whether these subsets showed differential enrichment for any of the 1,163 gene-sets (**Online Methods**). We found cluster A, related to the Q-wave, to be specifically enriched for 213 pathways related to structural abnormalities of the heart (**Fig. 3d**) such as “cardiac chamber development” and “dilated cardiomyopathy” (DCM). DCM is a severe clinical heart condition causing substantial morbidity and mortality^17^. The genetic variants of this cluster were indeed in close proximity to well known cardiomyopathy genes (e.g. *TTN, CAMK2D, OBSL1, BAG3* and *HSPB7/CLCNKA*)^18-20^.

To exploit the potential for obtaining new clinical insights of this resource, we focussed on a particular point of the Q-wave cluster that showed strong overlap with DCM risk. We found that rs2234962 in *BAG3* was among the most significant novel loci (P<5×10^-54^) and was specific for the Q-R upslope at −18ms from the R wave (**Fig. 4**, http://www.ecgenetics.org). This variant was previously identified in a GWAS of DCM^18-20^. Another newly identified ECG locus was rs1763604 near *HSPB7/CLCNKA* (P<5×10^-20^), also previously identified by a DCM GWAS and exhibiting the same ECG pattern (**Fig. 4**). Importantly, associations with this particular ECG morphologic feature persisted in individuals not diagnosed with DCM or other cardiac conditions **(Supplementary table 7)**. We hypothesized that this Q-R upslope feature (−18s from R wave) represents a biomarker for DCM risk. To further substantiate this hypothesis, we executed a Mendelian randomization analysis. Using up to 47 genetic variants, we observed that an increased Q-R slope is associated with increased risk of DCM (**Fig. 4c., Supplementary Table 8-9**). These findings remained consistent in several sensitivity analyses, across the UK Biobank and the independent MAGNet study cohort. Also exclusion of the 2 previously known DCM variants did not change our findings (**Fig. 4e.**). In addition to the known *BAG3* and *CLCNKA* loci, we identified 3 other DCM variants passing the bonferonni significant treshold (*P*<0.001): rs9909004 (*P*=3.5×10^-4^) in *PRKCA*, rs4685090 (*P*=4.2×10^-6^) near *LSM3* and rs60836537 (*P*=1.4×10^-4^) in *OBSCN* of this Q-R slope trait. The *PRKCA* locus has previously been found for dilated cardiomyopathy via adverse left ventricular modeling^21^. *LSM3* and neighboring genes have an unclear role in the heart. *OBSCN* encodes Obscurin, a signaling protein that plays a major role in the formation of new sacromeres during myofibril assembly and is harboring mutations causing DCM^22^. In total 13 of 47 variants were suggestively associated at *P*<0.05 with DCM, considerably more than expected by chance (*P*_binomial_ *=* 3.4 ×10^-7^. These analyses support this new hypothesis that the Q-R slope differences are linked to DCM risk. Further functional studies are important to understand the nature of these associations.

**Fig. 4.**
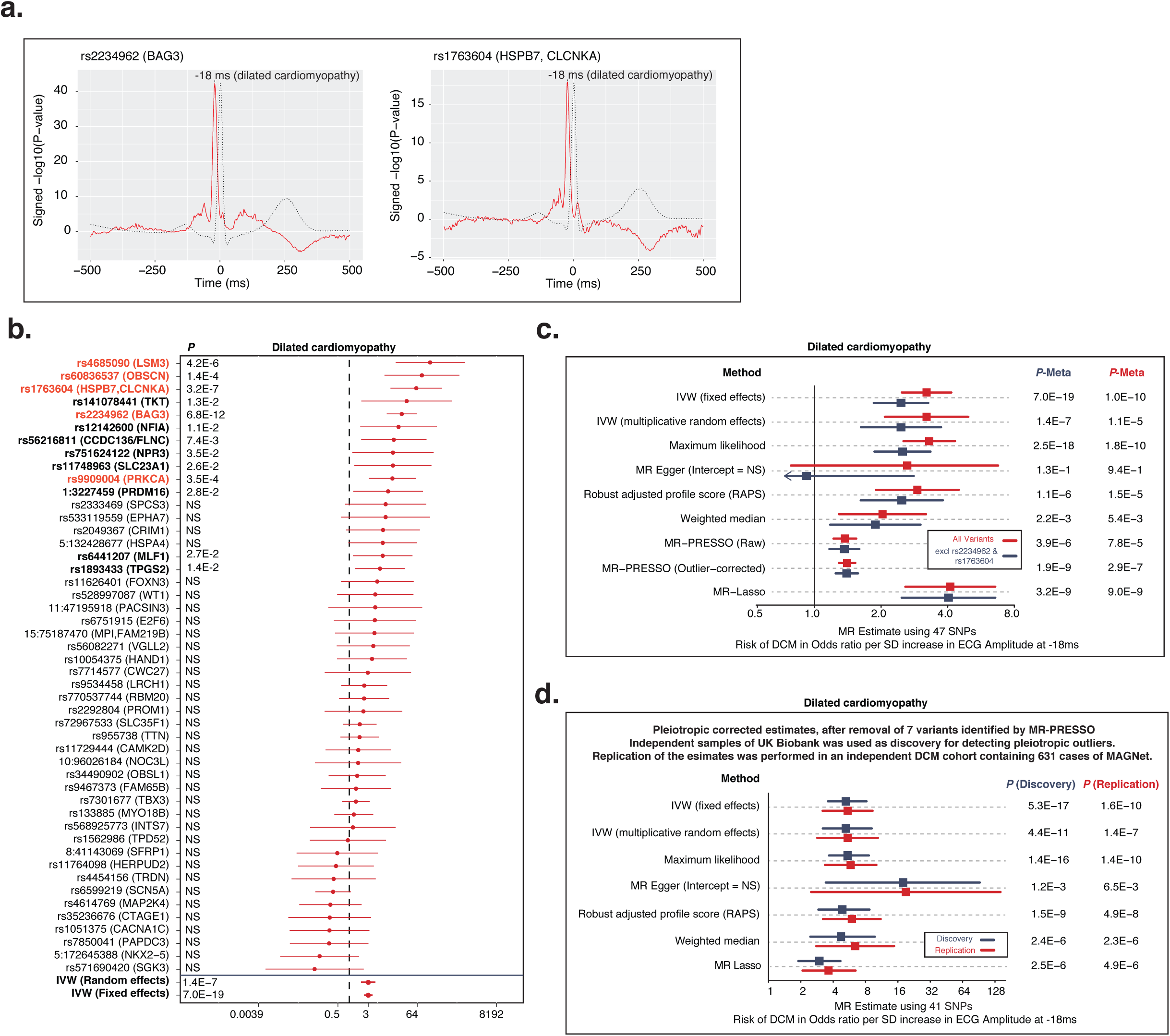
Mendelian randomization of dilated cardiomyopathy. **a**, The ECG morphology signature of previously identified loci associated with DCM. **b**, Individual variant effects on DCM for those significant for −18ms. **c**, Mendelian randomization estimates of the ECG at - 18ms on DCM **d.** Mendelian randomization estimates of the ECG at −18ms on DCM in the discovery (UK Biobank) and replication (MAGNet) cohort controlling for pleiotropic variants identified by MR-PRESSO.

Other examples of cardiac diseases in which the ECG is critical for clinical diagnosis include early repolarisation and the Brugada syndrome. The diagnosis of these conditions are based on strict criteria of passing a certain threshold on the ECG. We hypothesized that the biological underpinnings of these disorders do not adhere to a strict binary phenomenon, and that our understanding of these disorders can be enhanced by studying continuous traits as well. A previous GWAS of the Brugada syndome identified 3 independent genetic variants^23^, all of which were also among our 166 identified loci. Two of the 3 reported Brugada loci were independent signals in the *SCNSA* locus encoding the main cardiac sodium channel Na_v_1.5. The biology of the *SCNSA* locus is highly heterogenous as it has been linked to almost all other ECG traits^4,5,14,15,6-13^. To further dissect the complex genetic architecture of *SCNSA*, we performed a compehensive conditional analysis of this locus on the ECG (see **Supplementary Information**). We identified 10 independent variants in the *SCNSA* locus with an effect on one or more of the 500 data-points of the ECG-morphology (at P<5×10^-8^). These variants all coincide with cardiac enhancer marks (more than expected by chance, *P*_hypergeometric_=1×10^-7^). We found that that these 10 independent *SCNSA* variants have different morphological ECG signatures, possibly pointing towards different biological mechanisms and consequences for the cardiac conduction system (**Supplementary Data 4, Supplementary Table 10**).

Cardiovascular safety is a major concern for many drugs on the market and in development. Our data may contribute to early recognition of potential safety concerns. Drugs (off-) targeting ion channels are well established examples, including Na_v_1.5 (*SCNSA*), which is targeted by different drugs with cardiac safety concerns^24^. Another example are I_kr_ channels (*KCNH2*) targeted by class Ia anti-arrythmic drugs including quinidine and the class III anti-arrythmic drug sotalol, both known to prolong QT interval^25^, a feature also observed in our data (**Extended Fig. 1a**; http://www.ecgenetics.org), or drugs targeting Ca_v_3.1 (*CACNA1G*) known to affect the PR-interval^26^, a feature also observed in our data (**Extended Fig. 1b**; http://www.ecgenetics.org). We also anticipate that this work might be relevant to prioritze gene targets for therapy or more specifically monitor undesirable ECG effects, for example when prioritizing development of cancer immunotherapy.^27^

Although genetic influences on the ECG have long been established, these analyses have been based on isolated segments of the ECG. Here we provide the first high-dimensional analyses of the ECG that is not limited by the arbitrary human definition of particular ECG features. This novel resource provides a powerful tool to perform *in silico* analyses of genetic variants, genes and its relation to the ECG. We provided examples on how this resource opens up new avenues for studying cardiac physiology, as well as disease and drug development.

## Methods

### UK Biobank individuals

Participants were recruited with an age range of 40-69 years of age that registered with a general practitioner of the UK National Health Service (NHS). Between 2006-2010, in total 503,325 individuals were included. All study participants provided informed consent and the study was approved by the North West Multi-centre Research Ethics Committee. Detailed methods used by UK Biobank have been described elsewhere. The prevalence and incidence of cardiac conditions and events were captured by data collected at the Assessment Centre in-patient Health Episode Statistics (HES) download on September 10, 2017.

### Genotyping and imputation

The Wellcome Trust Centre for Human Genetics performed quality control before imputation and imputed to HRC v1.1 panel that was released on March 7th 2018. Quality control of samples and variants, and imputation was performed by the Wellcome Trust Centre for Human Genetics, as described in more detail elsewhere^28^. Individual sample outliers excluded based on heterozygosity and missingness were excluded, as well as those with gender discrepancies between the reported and inferred gender using X-chromosome heterozygosity test.

### Genetics and regression analyses

All of the genetic analyses in UK Biobank that are reported in this manuscript were adjusted for age, gender, BMI, height and the first 30 principal components (PCAs) to account for population stratification and genotyping array (Affymetrix UK Biobank Axiom® array or Affymetrix UK BiLEVE Axiom array). ECG variables were inverse rank normalized prior to the association analyses

Genome wide association analyses in UK Biobank were performed using BOLT-LMM v2.3beta2. BOLT-LMM fits a mixed linear model that accounts for the population structure and cryptic relatedness.^29^ For this, directly genotyped variants were used that passed quality control, and which were extracted from the imputed dataset to ensure 100% call rate, and pruned on linkage disequilibrium (first r^2^ < 0.05 and a second round of r^2^<0.045) to obtain roughly 400k variants across the genome. The genomic control lambda’s, heritability by BOLT-LMM, LD-Score intercepts and the attenuation ratio statistics are listed in **Supplementary Table 11**, these suggested no inflation due to non-polygenic signals ^30^. Regression analyses of genetic risk scores and individual genetic variant associations across the ECG-morphologies were performed with sandwich robust standard errors that were clustered by family to account for relatedness in STATA-MP v15. Families were inferred from the kinship matrix, clustering all 3rd degree relatives or higher together (kinship coefficient > 0.0442).

For the GWAS we focused on the ECG morphology unadjusted for R-R interval as it was more powerfull compared to the R-R adjusted morphology phenotype; higher heritability, captured the large majority of previously identified genetic variants of the ECG and was easiest to interpret. We performed 500 GWAS’s, one for each time-point. Using an eigenvalue-based measure^31^, we estimated that testing the 500 ECG data points resulted in effectively 79.9 independent tests and therefore used an alpha of 5×10^-8^/80 = 6.25×10^-10^ to indicate bonferroni corrected genome wide significance.

Normalized association profiles of variants were plotted as heatmaps where red colors indicate the most strongly associated effect at a time point and the blue color the most strongly associated effect in the opposite direction. To vizualize the similarities further on a 2-d plot, we used dimensionality reduction by t-sne on these normalized association profiles for all genome-wide variants, and k-means for clustering. This method only provides a global overview of the primary effects among variants.

### ECG morphology phenotypes and quality control

Three-lead exercise ECG data was provided by the UK Biobank as bulk in separate xml-files. To isolate R-waves, we employed the gQRS algorithm by George Moody, a public method previously found to be superior^32^, the construe algorithm was used to further refine the localization of R-waves ^33^. Individual ECG beats were processed and averaged^3^, which we refer to as the ‘ECG-morphology’ phenotype. Further source code of the methods and descriptions are available at https://github.com/niekverw/E-ECG.

Two entities of ECG morphology were constructed: The primary morphology trait was defined as the classical signal averaged electrocardiographic beat ^3^ that consisted of an averaged 1000ms window surrounding the R wave at a resolution of 500hz resulting in 500 averaged data points or ECG traits for each individual. As such, these beats were unadjusted for individual R-R intervals. The secondary ‘ECG morphology’ trait consisted of R-R intervals made of equal length (500 data points) so that the resulting averaged ECG beat was adjusted for each individual R-R interval.

Only information of the rest phase was used, as defined by the first 15 seconds of ECG assessment. From all 99,539 3-lead ECGs recorded in 96,567 participants, 77,190 ECGs contained full disclosure data nessesary to detect R wave. The R-wave is traditionally used as a reference point to detect all other points on the ECG, hence this should be sufficient to identify major changes ECG-beat, while also easy to understand and visualize.

Before signal averaging, ECG beats were quality controlled on the individual level. First, the 3-leads were averaged to create a single ECG signal vector. Then, individual beats containing excess noise were removed as described previously^34,35^, this was based on a moving standard deviation using a window-size of 3 data points under the assumption that ECG signals without noise have a moving standard deviation close to 0. Thirdly, individual beats were matched on a template and discarded if they were dissimilar based on a Pearson correlation function between each beat; beats with mean negative correlations and those that fell outside the standard 1.5 interquartile range rule were removed. This procedure was repeated until no outlier beats were left to exclude. At least 6 beats were required at any stage of the averaging process; otherwise the entire electrocardiogram was excluded from the analysis. We explored whether the averaging process was improved when accounting for the lag at which the cross-correlation function between each ECG beats shows its maximum, but this did not make a difference; suggesting that the R-wave detection was already sufficient. In total 67,440 ECGs of 66,240 individuals passed the individual level quality control.

Finally, to further detect and exclude abnormal ECGs on the population level, we calculated the standard deviation of the difference between each averaged ECG beat of an individual and the population-mean averaged ECG beat; outliers were discarded according to the standard 1.5 interquartile range rule per ECG-phenotype. Observations of the second follow-up visits were used when no baseline observation was available. This resulted in the inclusion of 63,706 individuals in the primary analysis (unadjusted R-R intervals). The 2,534 excluded individuals based on population level QC were much more likely to be diagnosed with a history of bundle branch block (OR=16.8, se= 3.1, z=15.41), cardiomyopathy (OR=8.8 se=2.3, z=8.58), myocardial infarction (OR=2.9, se=0.27, z=12.2), atrial fibrillation (OR=2.0, se=0.25, z=5.6), and heart failure (OR=7.3, se=1.2,z=12.55). For the secondary morphology phenotype (where we adjusted for R-R intervals), 65,183 individuals remained.

Another 3,750 individuals with a history of myocardial infarction, atrial fibrillation, heart failure, cardiomyopathy, bundle-branch-block or pacemaker implantation were excluded in sensitivity analyses and Mendelian randomization analyses of DCM.

Diagnoses in UK Biobank were defined as follows: myocardial infarction (ICD10: I21, I22, I23, I252; ICD9: 410, 412 or self reported myocardial infarction), atrial fibrillation (ICD10:I48; ICD9: 4273, OPCS-4: K621, K622, K623 or self reported atrial fibrillation or flutter), heart failure (ICD10: I50, I110, I130, I132; ICD9: 428 or self reported heart failure), cardiomyopathy (ICD10: I42, I43, O903, I255, O944; ICD9: 425 or self reported cardiomyopathy or hypertrophic cardiomyopathy), dilated cardiomyopathy (ICD10:I42.0 or ICD9: 4254), bundle-branch-block (ICD10: I447, I451 or ICD9: 4263, 4264) or pacemaker implantation (OPCS-4: K59, K60, K61 or self reported pacemaker or defibrillator insertion). Diagnoses in UK Biobank were extracted using https://github.com/niekverw/ukpheno.

### Polygenic scores of ECG-traits and known genetic variants

Polygenic risk scores were created following an additive model for atrial fibrillation, QT-interval, QRS duration, PR-interval, QRS-voltage traits (12-lead sum area, cornel area and Sokolow-Lyon area) and heart rate separately (**Supplementary Table 12**), as previously described^36^. In short, the number of alleles for each individual (0, 1 or 2) was summed after multiplying the alleles with the previously reported effect size of the variant-trait association. Effect sizes estimated in UK Biobank data were avoided to reduce potential overestimation. If multiple effect sizes were available, those estimated in the largest sample size were used (eg, the combined replication and discovery phase). Single-nucleotide polymorphisms were excluded if they were missing in UK Biobank data. In instances where multiple correlated variants in the same locus were reported for the same trait, we used only independent variants that were selected by the slinkage disequilibrium clumping procedure (at r^2^ < 0.01) implemented in PLINK version 1.9.

### Pathway analyses and candidate genes

DEPICT was employed to discovery pathways, tissues and genes underlying gwas-loci of the electrocardiographic morphology phenotype. Please see Pers *et al.* for a detailed description of the method^16^. All independently associated variants (r2>0.005) passing a traditional genome wide significance treshold P < 5 × 10^-8^ were used as input to the DEPICT framwork. DEPICT was run using default settings. Because DEPICT uses 1000 Genomes as a reference panel which does not include certain UK Biobank specific varianst, we also included all variants in LD (r^2^>0.8) with the input-variants to ensure 100% coverage of the signal. Pairwise correlations across all significantly enriched pathways (false discovery rate > 0.5%) were computed and vizualized using the Gephi software (www.gephi.org) after filtering out edges with correlations lower than 0.5.

For the differential pathway analysis, we repeated the DEPICT pathway analysis but excluded variants that were of interest (variants-of-interest). We calculated the reduction in pathway enrichment by substracting the −log_10_(*P*) for enrichment before and after excluding variants-of-interest. A null distribution was created by repeating the DEPICT run 100 times on the same set of variants but each time excluding a matched number of random variants not belonging to the variants-of-interest. If all 100 null runs indicated that the reduction in significance for a given pathway was more using the variants-of-interest than for the null runs (indicating *P*_differential_<0.01), the pathway was considered to be differentially enriched for the variants-of-interest. Particularly, we were interested whether different biological pathways were active at different ECG intervals. To test this, we used the clusters of variants that were identified in the unbiased clustering aproach by t-sne and k-means as ‘variants-of-interest’ (see *Genetics and association testing*); which grouped variants by their primary effect on the different ECG wave intervals in an unbiased manner.

Nearest gene or any gene within 10kb of the lead variants were used to annotate candidate causal genes. We also searched for coding variants in LD with the identified variants (r^2^>0.8) to further prioritize candidate causal genes. An additional line of evidence for candidate causal genes was performed by DEPICT, taking into account gene-gene similarities across loci^16^. We specifically did not incorporate annotations from datasets such as Hi-C and eQTL, as we strongly believe these may not be informative for cardiac conduction biology and need targeted experiments on a per locus basis to interpret them ^37,38^.

### Dilated cardiomyopathy

The association between genetic variants and risk of dilated cardiomyopathy (DCM) was tested in case-control GWASes of UK Biobank and MAGNet (Myocardial Applied Genomics Network) and combined via inverse-variance weighted meta-analysis to increase power totalling 1,375 cases and 241,325 controls.

In the UK Biobank cohort individuals with and without DCM were selected to be independent of those taking part in the ECG measurements. 744 DCM cases were identified according to ICD10 and ICD9 codes (I42.0 and 4254), 239,729 control individuals were not diagnosed with DCM and had no family history of heart disease. Association testing was performed as described under *‘Genetic and association testing’*, coefficients were re-scaled to log odds.

For the MAGNet study, 631 subjects with dilated cardiomyopathy were recruited; patients were included in the study if they were diagnosed with heart failure with reduced ejection fraction (< 40%) in the absence of hypertension, primary valvular disease, or coronary artery disease from the University of Pennsylvania Health System; 1,596 controls were recruited from the same center who had no history of heart disease. All subjects provided written informed consent. Genotyping was performed on the Illumina OmniExpressExome Array and imputed using the 1000 Genomes Project with Minimac. Genotypes between cases and controls were compared using an additive genetic model adjusting for gender and two principal components of race using SNPtest; there was no genomic inflation (1.0335).

### Mendelian randomization

Lead variants associated at *P* < 5 × 10^-8^ were used as instrumental variables in the MR. Enrichment of low P values (*P* < 0.05) among variants was calculated using a binomial distribution. Mendelian randomization analyses included the inverse-variance-weighted fixed-effects, random-effects meta analyses and weighted median. Heterogeneity was assessed through I^2^ index and Cochran’s Q. Pleiotropy analyses included the MR-Egger intercept, MR-RAPS and MR-LASSO estimates^39^. MR pleiotropy residual sum and outlier (MR-PRESSO)^40^ was used as MR analysis to detect and remove pleiotropic variants. MR-PRESSO accounts for pleiotropic effects of the genetic variants comparing the observed distance of all the variants to the regression line (residual sum of squares) to the expected distance under the null hypothesis of no horizontal pleiotropy. MR-PRESSO is based on the assumption that at least 50% of the variants are valid instruments.

For the Mendelian randomization of DCM, variant effects on the ECG morphology at −18 ms were estimated using only individuals without a history of cardiac disease. We performed an additional analysis to explore the impact of pleitropic signals by conducting a 2-stage Mendelian randomization using a discovery and replication stage. First we assessed the causal relationsip in UK Biobank and identified pleitropic outliers by MR-PRESSO. Secondly, we replicated this analysis by repeating the MR in an independent cohort of 631 DCM patients from MAGNet. In this second stage, we excluded the variants identified by MR-PRESSO in the first stage, hence correcting for pleiotropic effects in an unbiased manner.

## Data availability

Genome-wide summary statistics can be queried and downloaded from www.ecgenetics.org. Further datasets generated and/or analysed during this study are available from the corresponding authors on reasonable request.

### Acknowledgements

This research has been conducted using the UK Biobank Resource under Application Number 12010. We thank Ruben N. Eppinga, MD, M. Yldau van der Ende, BSc, and Yanick Hagemeijer, MSc, University of Groningen, University Medical Center Groningen, Department of Cardiology, for their contributions to the extraction and processing of data in the UK Biobank. None of the mentioned contributors received compensation, except for their employment at the University Medical Center Groningen.

This work was supported by NWO VENI grant 016.186.125 and NIH grants HL105993 and HL088577. The funders had no role in the design and conduct of the study; collection, management, analysis and interpretation of the data; preparation, review or approval of the manuscript; or decision to submit the manuscript for publication.

---

**Extended Data Fig 1. Examples of loci relevant for known drugs targeting ion channels, illustrating the predicted drug effect.**

## References

1. Das, M. K. & Zipes, D. P. Electrocardiography of arrhythmias: a comprehensive review: a companion to Cardiac electrophysiology: from cell to bedside. (Elsevier/Saunders, 2012).

2. AlGhatrif, M. & Lindsay, J. A brief review: history to understand fundamentals of electrocardiography. *J*. community Hosp. Intern. Med. Perspect. 2, (2012).

3. Simson, M. B. Use of signals in the terminal QRS complex to identify patients with ventricular tachycardia after myocardial infarction. Circulation 64, 235-42 (1981).

4. Verweij, N. et al. Twenty-eight genetic loci associated with ST-T-wave amplitudes of the electrocardiogram. Hum. Mol. Genet. 25, 2093-2103 (2016).

5. van der Harst, P. et al. 52 Genetic Loci Influencing Myocardial Mass. *J*. Am. Coll. Cardiol. 68, 1435–l1448 (2016).

6. Eppinga, R. N. et al. Identification of genomic loci associated with resting heart rate and shared genetic predictors with all-cause mortality. Nat. Genet. 48, 1557–1563 (2016).

7. van den Berg, M. E. et al. Discovery of novel heart rate-associated loci using the Exome Chip. Hum. Mol. Genet. 26, 2346–2363 (2017).

8. Arking, D. E. et al. Genetic association study of QT interval highlights role for calcium signaling pathways in myocardial repolarization. Nat. Genet. 46, 826–836 (2014).

9. Bihlmeyer, N. A. et al. ExomeChip-Wide Analysis of 95 626 Individuals Identifies 10 Novel Loci Associated With QT and JT Intervals. Circ Genom Precis Med 11, e001758 (2018).

10. van Setten, J. et al. PR interval genome-wide association meta-analysis identifies 50 loci associated with atrial and atrioventricular electrical activity. Nat. Commun. 9, 2904 (2018).

11. Roselli, C. et al. Multi-ethnic genome-wide association study for atrial fibrillation. Nat. Genet. (2018). doi:10.1038/s41588-018-0133-9

12. Nielsen, J. B. et al. Biobank-driven genomic discovery yields new insight into atrial fibrillation biology. Nat. Genet. 50, 1234–1239 (2018).

13. Prins, B. P. et al. Exome-chip meta-analysis identifies novel loci associated with cardiac conduction, including ADAMTS6. Genome Biol. 19, 87 (2018).

14. Christophersen, I. E. et al. Fifteen Genetic Loci Associated With the Electrocardiographic P Wave. Circ. Cardiovasc. Genet. 10, (2017).

15. Verweij, N. et al. Genetic Determinants of P Wave Duration and PR Segment. Circ. Cardiovasc. Genet. 7, 475–481 (2014).

16. Pers, T. H. et al. Biological interpretation of genome-wide association studies using predicted gene functions. Nat. Commun. 6, 5890 (2015).

17. McKenna, W. J., Maron, B. J. & Thiene, G. Classification, Epidemiology, and Global Burden of Cardiomyopathies. Circ. Res. 121, 722–730 (2017).

18. Meder, B. et al. A genome-wide association study identifies 6p21 as novel risk locus for dilated cardiomyopathy. Eur. Heart J. 35, 1069–1077 (2014).

19. Stark, K. et al. Genetic Association Study Identifies HSPB7 as a Risk Gene for Idiopathic Dilated Cardiomyopathy. PLoS Genet. 6, e1001167 (2010).

20. Villard, E. et al. A genome-wide association study identifies two loci associated with heart failure due to dilated cardiomyopathy. Eur. Heart J. 32, 1065–1076 (2011).

21. Hu, R. et al. Genetic Reduction in Left Ventricular Protein Kinase C-a and Adverse Ventricular Remodeling in Human Subjects. Circ. Genomic Precis. Med. 11, e001901 (2018).

22. Makarenko, I. et al. Passive Stiffness Changes Caused by Upregulation of Compliant Titin Isoforms in Human Dilated Cardiomyopathy Hearts. Circ. Res. 95, 708–716 (2004).

23. Bezzina, C. R. et al. Common variants at SCN5A-SCN10A and HEY2 are associated with Brugada syndrome, a rare disease with high risk of sudden cardiac death. Nat. Genet. 45, 1044–1049 (2013).

24. Erdemli, G. et al. Cardiac Safety Implications of hNav1.5 Blockade and a Framework for Pre-Clinical Evaluation. Front. Pharmacol. 3, 6 (2012).

25. Extramiana, F. et al. The Time Course of New T-Wave ECG Descriptors Following Single- and Double-Dose Administration of Sotalol in Healthy Subjects. Ann. Noninvasive Electrocardiol. 15, 26–35 (2010).

26. Oparil, S. et al. Dose-Response Characteristics of Mibefradil, a Novel Calcium Antagonist, in the Treatment of Essential Hypertension. Am. J. Hypertens. 10, 735–742 (1997).

27. Patel, S. J. et al. Identification of essential genes for cancer immunotherapy. Nature 548, 537–542 (2017).

28. Bycroft, C. et al. The UK Biobank resource with deep phenotyping and genomic data. Nature 562, 203–209 (2018).

29. Loh, P.-R. et al. Efficient Bayesian mixed-model analysis increases association power in large cohorts. Nat. Genet. 47, 284–90 (2015).

30. Loh, P.-R., Kichaev, G., Gazal, S., Schoech, A. P. & Price, A. L. Mixed-model association for biobank-scale datasets. Nat. Genet. 50, 906–908 (2018).

31. Galwey, N. W. A new measure of the effective number of tests, a practical tool for comparing families of non-independent significance tests. Genet. Epidemiol. 33, 559–568 (2009).

32. Computing in Cardiology Conference 2Ol4 (CinC): September 7-lO, 2Ol4, Cambridge, Massachusetts, USA.

33. Teijeiro, T., Felix, P., Presedo, J. & Castro, D. Heartbeat Classification Using Abstract Features From the Abductive Interpretation of the ECG. IEEE J. Biomed. Heal. Informatics 22, 409–420 (2018).

34. Verweij, N., Van De Vegte, Y. J. & Van Der Harst, P. Genetic study links components of the autonomous nervous system to heart-rate profile during exercise. Nat. Commun. 9, (2018).

35. van de Vegte, Y. J., van der Harst, P. & Verweij, N. Heart rate recovery 10 seconds after cessation of exercise predicts death. J. Am. Heart Assoc. 7, (2018).

36. Verweij, N., Eppinga, R. N., Hagemeijer, Y. & Van Der Harst, P. Identification of 15 novel risk loci for coronary artery disease and genetic risk of recurrent events, atrial fibrillation and heart failure. Sci. Rep. 7, (2017).

37. Wang, X. & Goldstein, D. B. Enhancer redundancy predicts gene pathogenicity and informs complex disease gene discovery. bioRxiv 459123 (2018). doi:10.1101/459123

38. Stacey, D. et al. ProGeM: a framework for the prioritization of candidate causal genes at molecular quantitative trait loci. Nucleic Acids Res. 47, e3–e3 (2019).

39. Slob, E. A. W. & Burgess, S. A Comparison Of Robust Mendelian Randomization Methods Using Summary Data. bioRxiv 577940 (2019). doi:10.1101/577940

40. Verbanck, M., Chen, C.-Y., Neale, B. & Do, R. Detection of widespread horizontal pleiotropy in causal relationships inferred from Mendelian randomization between complex traits and diseases. Nat. Genet. 50, 693–698 (2018).

