## Supplementary figures and images for "The genetic makeup of the electrocardiogram"

### Supplemental Data 1

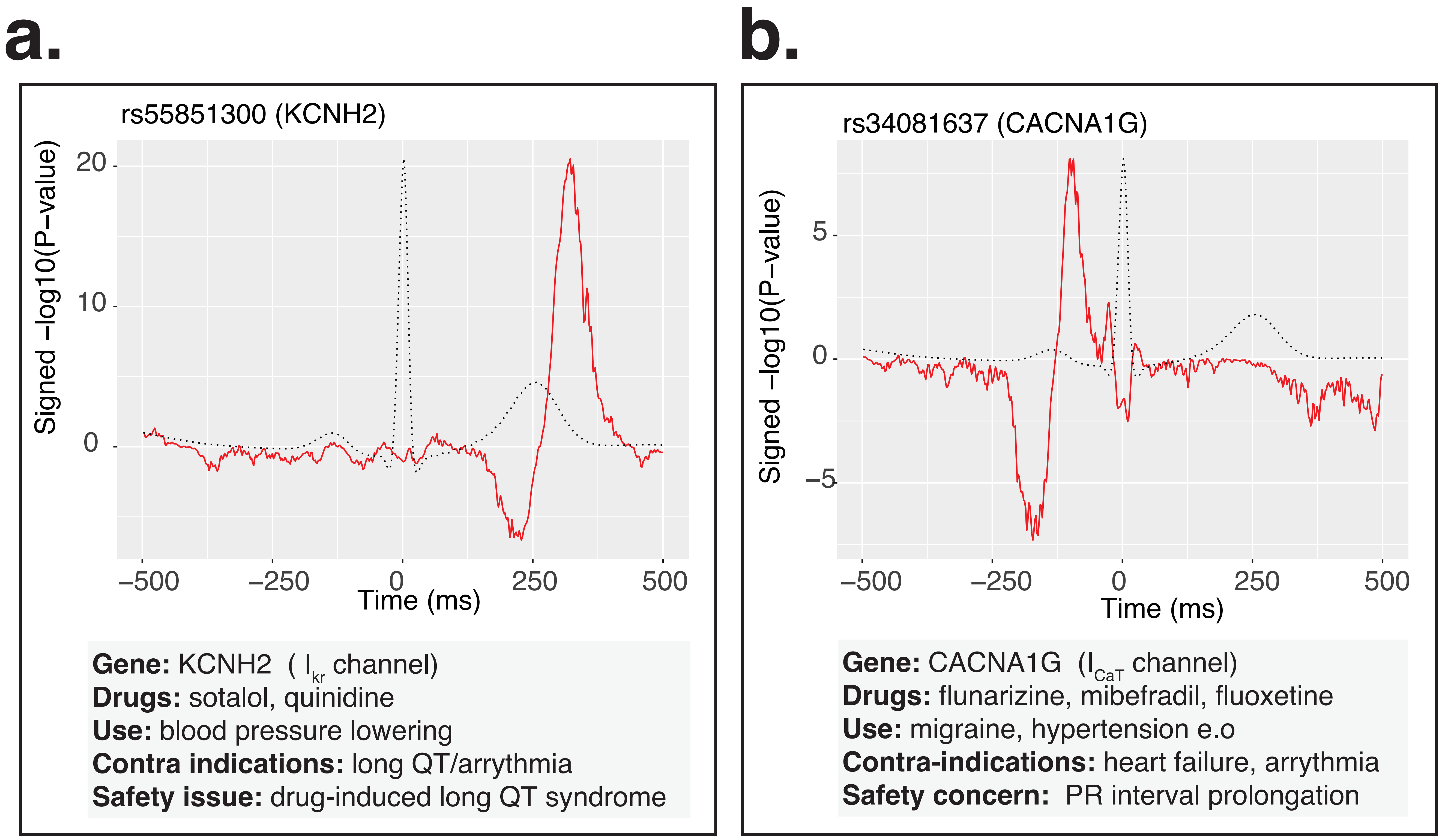
