## Supplementary Data 4 for "The genetic makeup of the electrocardiogram"

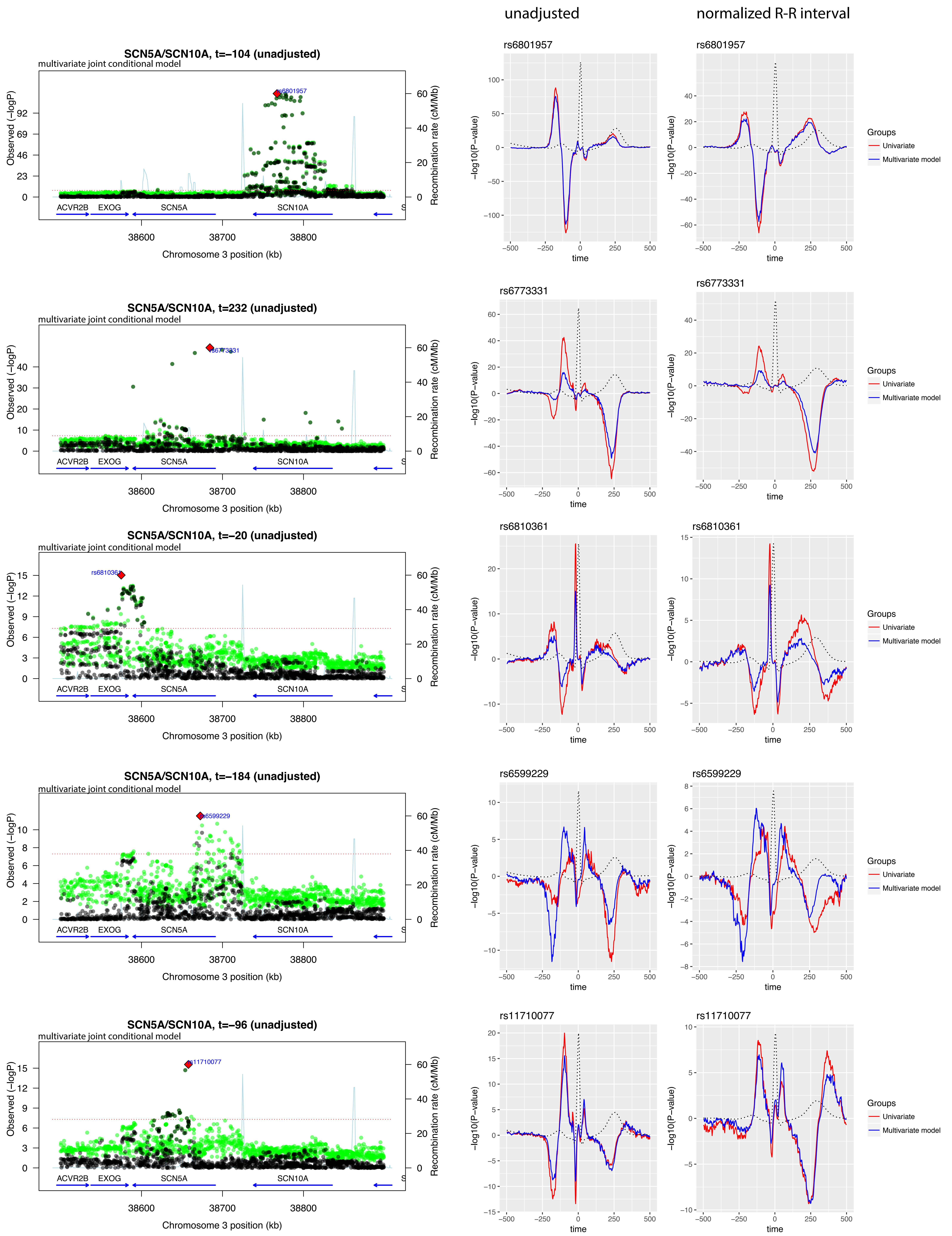


Supplementary Figure. Finemapping the SCN5A locus. The regional plots are for each of the 10 independent signals, conditioned for the other 9 variants. black points depict the ECG-time for which the variant was most significant, green points are the minimum P values across the ECG morphology phenotype. On the right of the regional plots are the associations of each variant plotted on the ECG morphology (unadjusted and adjusted for RR interval). The ed lnine indicates the association statistic in the univariate model (no snps included in the model), the blue lines depicts the -logP value from the multivariate joint conditional model containing respectively all other 9 genetic variants.

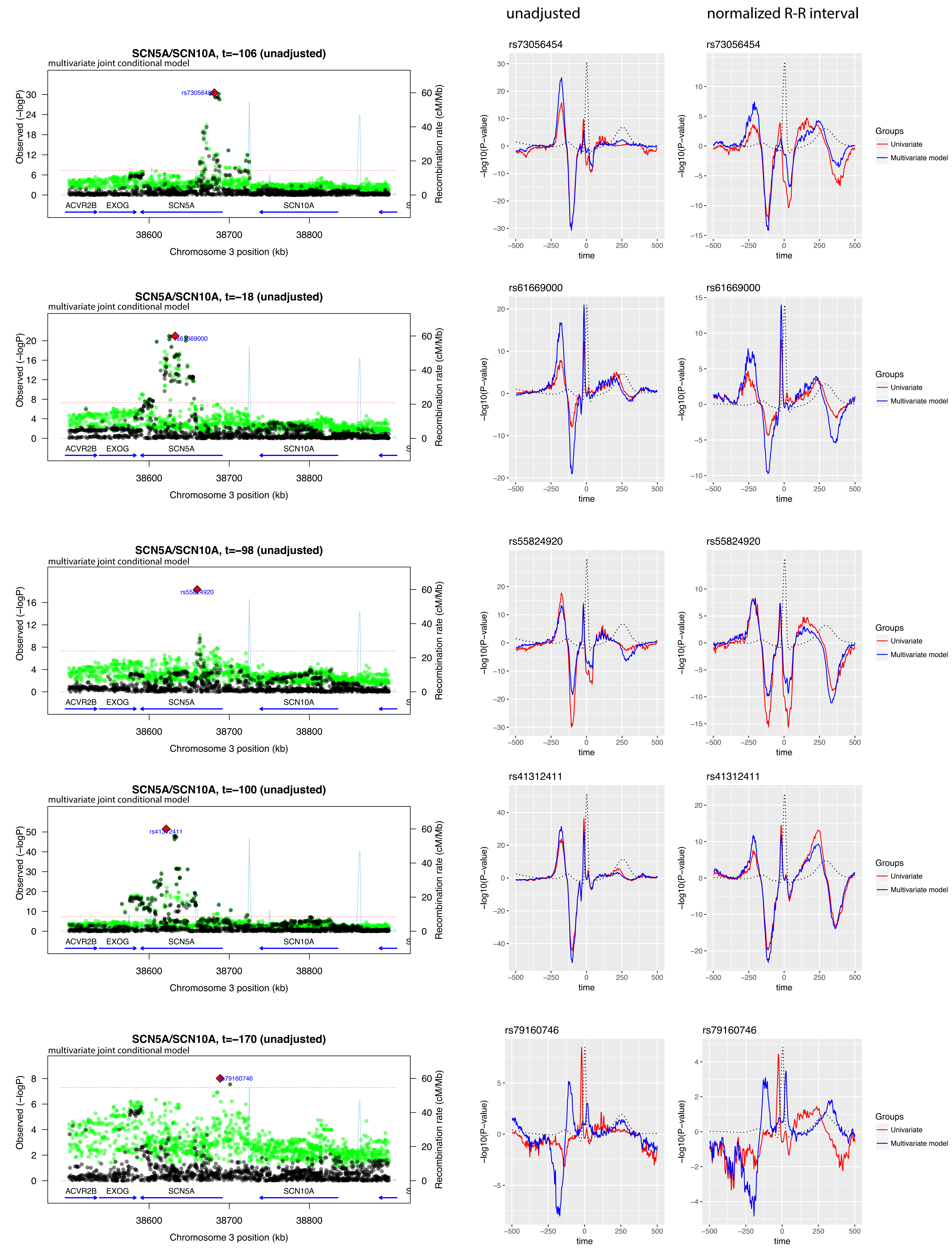


Supplementary Figure.  Continued

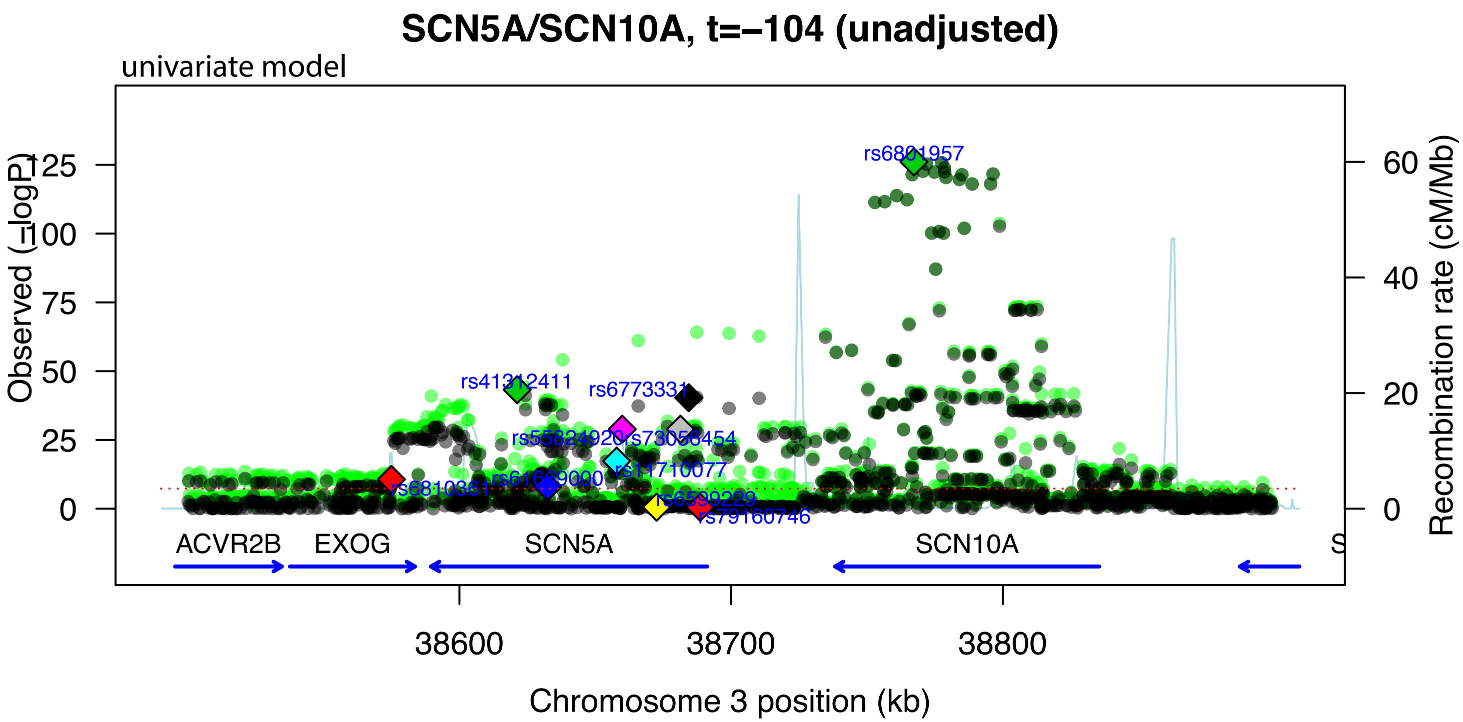

SCN5A/SCN10A, t=−104 (unadjusted)
univariate model
Observed (−logP)
125
100
75
50
25
0
Recombination rate (cM/Mb)
60
40
20
0
rs6801957
rs41312411
rs6773331
rs55824920
rs73058454
rs11710077
rs61669000
rs6810361
rs6599229
rs79160746
ACVR2B
EXOG
SCN5A
SCN10A
38600
38700
38800
Chromosome 3 position (kb)
