## supplementary Materials for "The genetic makeup of the electrocardiogram"

Niek Verweij et al.

**Exhaustive joint conditional analysis of the SCN5A-SCN10A locus**

The SCN5A locus has been reported in almost all cardiac conduction GWAS with various independent lead SNPs, it harbors many mutations causing different types of arrhythmias, and it encodes an essential sodium channel $Na_v1.5$ that is a common target of anti-arrhythmic drugs. To further increase our understanding of complex genetics at SCN5A, we performed an exhaustive joint conditional analysis across all ECG datapoints. Finemapping this locus and understanding the consequences of the multiple genetic variants at play may improve our understanding of this gene and cardiac disease pathophysiology.

We performed an in-depth conditional analysis of this locus using information of the entire ECG using a deterministic approach. For this, a stepwise conditional analysis was conducted followed by the full model containing all of the identified SNPs P< x $10^{-8}$; after this the stepwise conditional analysis was redone with as starting point the SNPs that were also the lead SNP in the full model. This was repeated until all SNPs in the model were the lead associated SNP in the full model.

The first stepwise model including all 1309 variants between 38500000 and 38900000bp on chromosome 3 identified 11 genetic variants. After 3 iterations, a final model was constructed of 10 genetic variants that were independently associated and also the top-associated variants across all ECG association analyses. Please see **Supplementary Table 11** for more information on the exact configurations that were tested and **Supplementary Figure 11** for the conditional regional association plots and impact on ECG morphology. While the genetic signal was clearly separated out by this approach, the phenotypic associations were not very much changed compared to the unconditioned analyses (**Supplementary Figure 11**).

Notably, all of the 10 genetic variants were located in a cardiac enhancer (https://pubs.broadinstitute.org/mammals/haploreg). When summarizing the overlap of all variants in the *SCN5A* region, only 21% overlapped with a cardiac enhancer, this 10 out of 10 genetic variants represents a considerable enrichment P = 1 x $10^{-7}$ on the single locus level (hypergeometric test). This suggests that the causal common variants at *SCN5A* are likely to be of regulatory nature. And although these variants may not be considered the causal variants underlying the ECG, this does provide some extra support for the hypothesis that these variants may be functional.

Previously identified genetic variants for Brugada syndrome at rs10428132 and rs11708996 were also found back as independent signals in this analysis. Rs10428132 was the genetic variant with the lowest P values overall and which was fine-mapped to rs6801957; rs11708996 was in perfect LD with rs41312411, highlighting that this approach is able to capture disease relevant genetic variants. Both variants had a strong effect on the P-wave interval, whereas Brugada is characterized by ST abnormalities, the syndrome is also known to present a concealed abnormal atrial, P-wave, phenotype [1] .

rs41312411 also has a specific effect on the beginning of the QRS complex, which seems to affect Q-R duration and/or Q amplitude.

**References**

1. Conte, G. *et al.* Concealed abnormal atrial phenotype in patients with Brugada syndrome and no history of atrial fibrillation. *Int. J. Cardiol.* **253,** 66–70 (2018).
